# Optimized Protocol for Cyanobacterial 16S rRNA Analysis in Danube Delta Lakes

**DOI:** 10.1101/2021.04.23.441086

**Authors:** Maria Iasmina Moza, Carmen Postolache

## Abstract

Molecular biology protocols have been more and more accessible to researchers for ecological investigations, however, these protocols always require optimization steps for the analysis of specific types of samples. The purpose of this study was to optimize a molecular protocol for the analysis of cyanobacterial 16S rRNA in Danube Delta shallows lakes. In this regard, several commercial DNA extraction kits were tested in comparison with potassium ethyl xanthogenate extraction method on different matrices. The obtained DNA was further used for 16S rRNA PCR optimization. Finally, an optimized protocol is proposed for the molecular analysis of cyanobacteria group in freshwater samples. The best DNA extraction method was the potassium xanthogenate extraction from dried cyanobacterial biomass. A dynamic in total genomic eDNA was observed, reflecting the seasonal difference in phytoplankton biomass from the studied lakes. The PCR protocol optimized by us can be successfully applied for the identification of a broad range of cyanobacterial genetic markers.

## 1. Introduction

Cyanobacteria are an ancient group of autotrophic bacteria living both in freshwater and marine environments and constituting an important component of the primary producers (Moore et al. 2019, Whitton and Potts 2007). Mass populations of toxic cyanobacteria represent a global phenomenon and the recent recognition that incidences of blooms may increase significantly under climate change serves to reinforce further the seriousness of the potential risks to human health (Paerl and Huisman 2008). In recent years the increased temperatures triggered a higher frequency of cyanobacteria blooming, including species with potential to release toxins also in the Danube Delta Biosphere Reserve (Török 2005), and thus accentuated the health risk hazards. There is a high diversity of aquatic ecosystems here, and consequently, a high diversity and variability of phytoplankton community, that makes difficult to predict the occurrence of toxic cyanobacteria blooms only by using the classic methods (e.g. microscopy or fluorometry). For this reason, molecular analyses focusing on the identification of genes able to release toxins during the blooms may complement the classic methods and contribute to predict a certain “toxic bloom” pattern in order to reduce casualties.

Since the majority of microorganisms are uncultivable in the laboratory, DNA isolation and PCR provide a powerful tool for studying microorganisms directly from environmental samples (Yilmaz et al. 2009). These techniques are currently used in microbial ecology (Martins et al. 2011) for the identification of a broad range of cyanobacterial genetic markers and for the quantification of structural and functional properties of cyanobacterial communities in both field and laboratory conditions (Albrecht et al. 2017, Moreira et al. 2013). However, the protocol for DNA extraction requires optimization in the case of environmental samples, for particular matrices, and for certain taxonomic groups respectively (Abdel-Latif and Osman 2017, Akkak et al. 2008, Chandraa et al. 2010, Dobhal et al. 2014, Drábková et al. 2002, Mirmomeni et al. 2010, Morin et al. 2010, Nair et al. 2014, Verbylaite et al. 2010).

For cyanobacteria DNA extraction, adapted protocols are needed even for each group apart (Billi et al. 1998) since their cells walls contain large amounts of cellulose, pectins, murein and xylose (Kaczyńska et al. 2013) which interfere with the cell lysis and DNA isolation leading to only small amounts of DNA, also contaminated. Moreover, cyanobacteria form a mucous envelope, which protects the cells against various environmental factors (Kaczyńska et al. 2013). Particularly, benthic forms,- can produce protective sheaths or mucilage (Gaget et al. 2017) that harden the analysis since they interfere with DNA extraction. For the Danube Delta we reported plenty of filamentous, but also colony forming species (Moza et al. 2021). Most of them have enormous quantities of muco-polysaccharides that make the DNA extraction very challenging (Tiam et al. 2019). Quality and quantity of extracted DNA can be then tested using two methods: spectrophotometer quantification, PCR and electrophoresis on agarose gel (Chandraa et al. 2010).

This study represents the first attempt focusing on culture-independent studies of cyanobacteria from Danube Delta shallow lakes. Therefore, we aimed to: (1) find the best methods for extraction of high-quality DNA using different preservation protocols and several commercial kits and the lab made potassium ethyl xanthogenate extraction buffer (XS buffer) and (2) to optimize a PCR protocol for 16S ribosomal RNA gene amplification.

## 2. Materials and methods

### 2.1. Study sites

The Danube Delta is a Biosphere Reserve (DDBR) located at 45°0’N latitude and 29°0’E longitude in the eastern part of Romania and comprises more than 400 lakes, well interconnected by a complex network of natural and man-made channels as well as by river branches (Coops et al. 2008, Romanescu 2005). We sampled a number of lakes (Fig. 1 and Table 1) belonging to all four lake complexes (LCs) of DDBR, namely: Şontea-Furtuna (LC 1), Isac-Gorgova (LC 2), Matiţa-Merhei (LC 3), forming the fluvial delta, and Roşu-Puiu (LC 4) part of the maritime delta (Oosterberg et al. 2000). The heterogeneity of the LCs also can be highlighted by the water level which can be a surrogate for the hydrological regime (Irimuş 2006) and it was shown to influence not only the water volume sampled but as well the ecological state of the lakes (Moza et al. 2021). Water level can be considered, at the same time, driver of the cyanobacteria community distribution in shallow lakes (Moza et al. 2021). The water level in the DDBR lakes usually increases from West to East, following the water flow direction of the Sulina main branch. Figure 2 shows that the maximum water level in Danube Delta lakes during the dry period, when no pressures acted on the water regime (e.g. flooding’s) was registered in LC 4 (Roşu-Puiu) the most eastern complex, forming the maritime delta. A detailed limnological description of the lakes was already done in previous papers (Enache et al. 2019, Fontana et al. 2018). The sampling campaigns were carried out preliminary in October 2012 (12 lakes) and in spring (May), summer (July) and autumn (September) in 2013 (26 lakes) in order to include the seasonal dynamics of the cyanobacteria communities.

**Figure 1.**
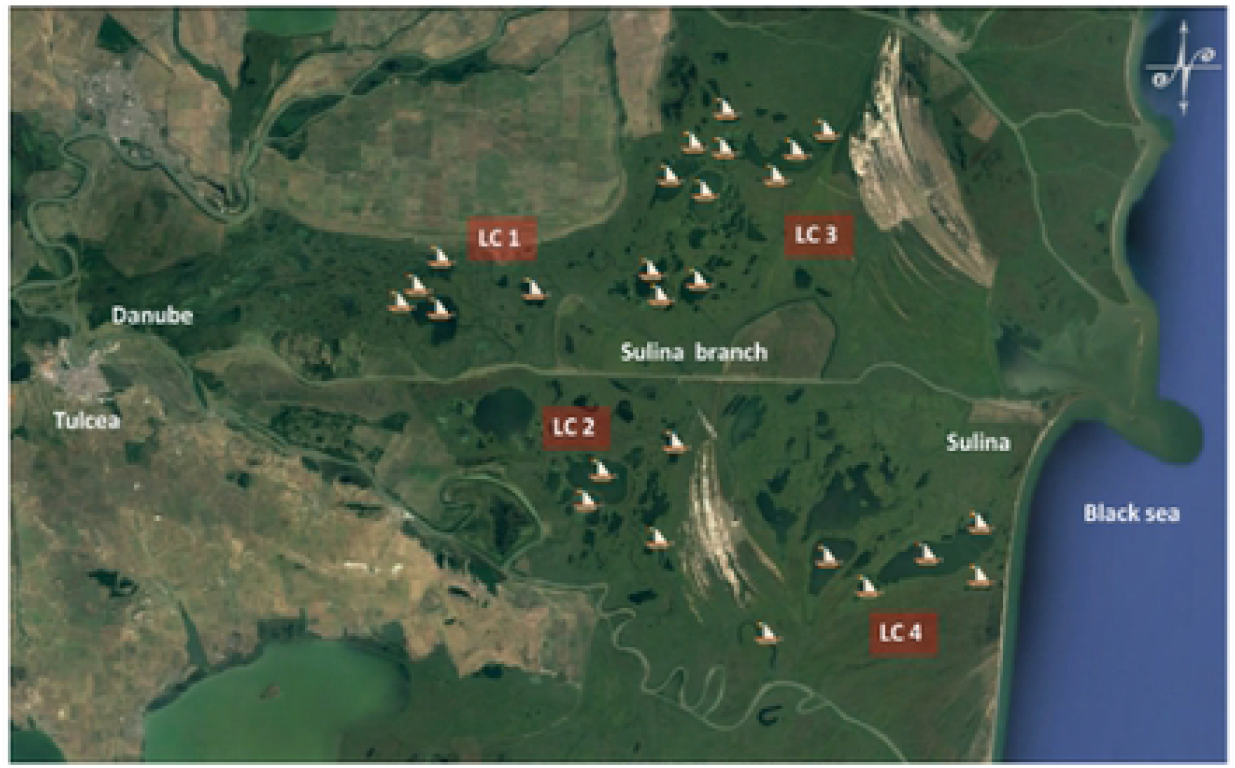
Danube Delta map (obtained from Google Earth Pro in 29.04.2020) with lakes (indicated with boats) and lake complexes (LC): Şontea-Furtuna (LC 1), Isac-Gorgova (LC 2), Matiţa-Merhei (LC 3) and Roşu-Puiu (LC 4) sampled during our survey in 2013

**Table 1.**
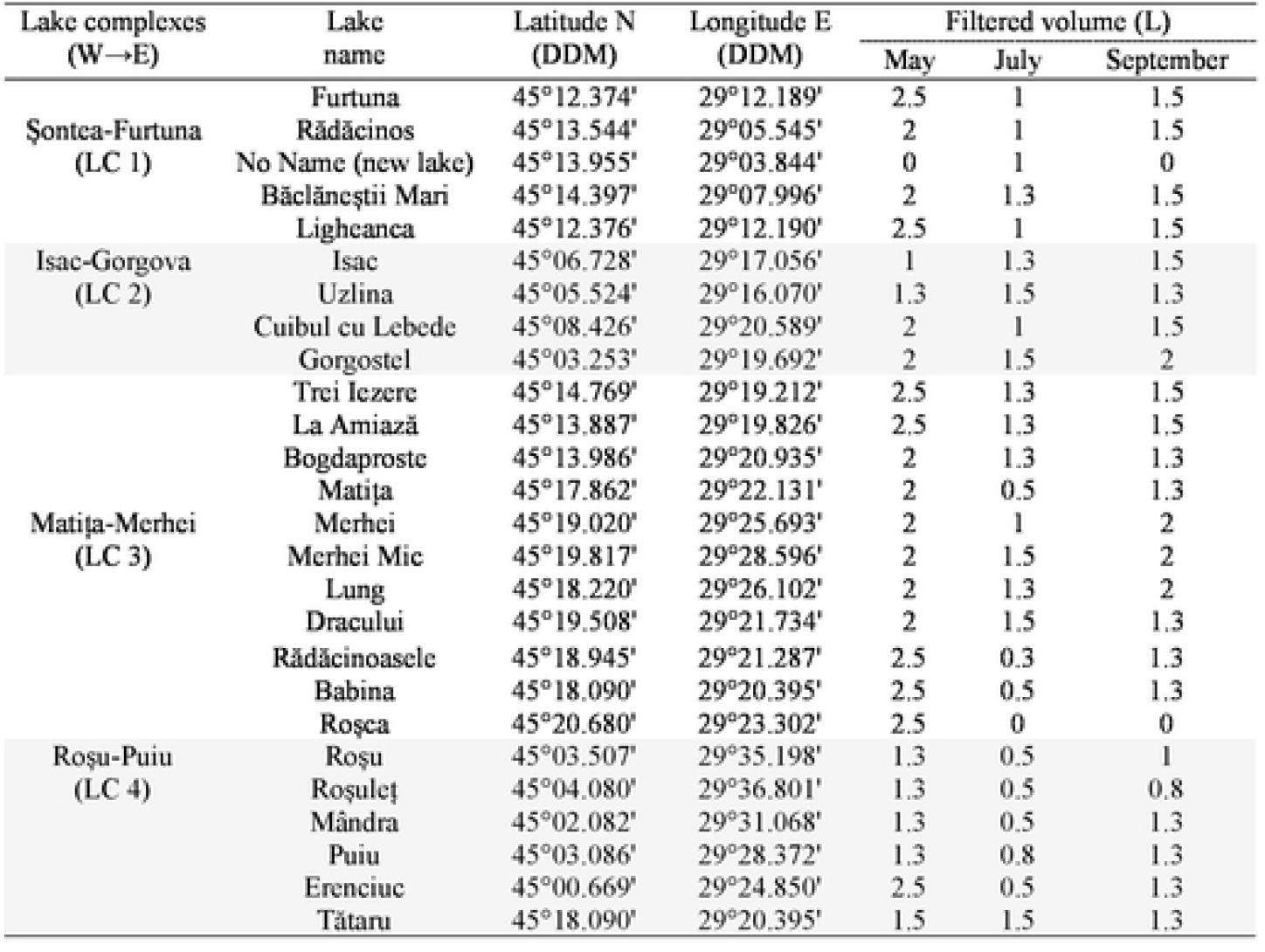
Lakes sampled from Danube Delta Biosphere Reserve in 2013 with their GPS coordinates of the sampling points (center of each lake) and the filtered water volume used for eDNA; lakes are listed from West to East along Sulina main branch

**Figure 2.**
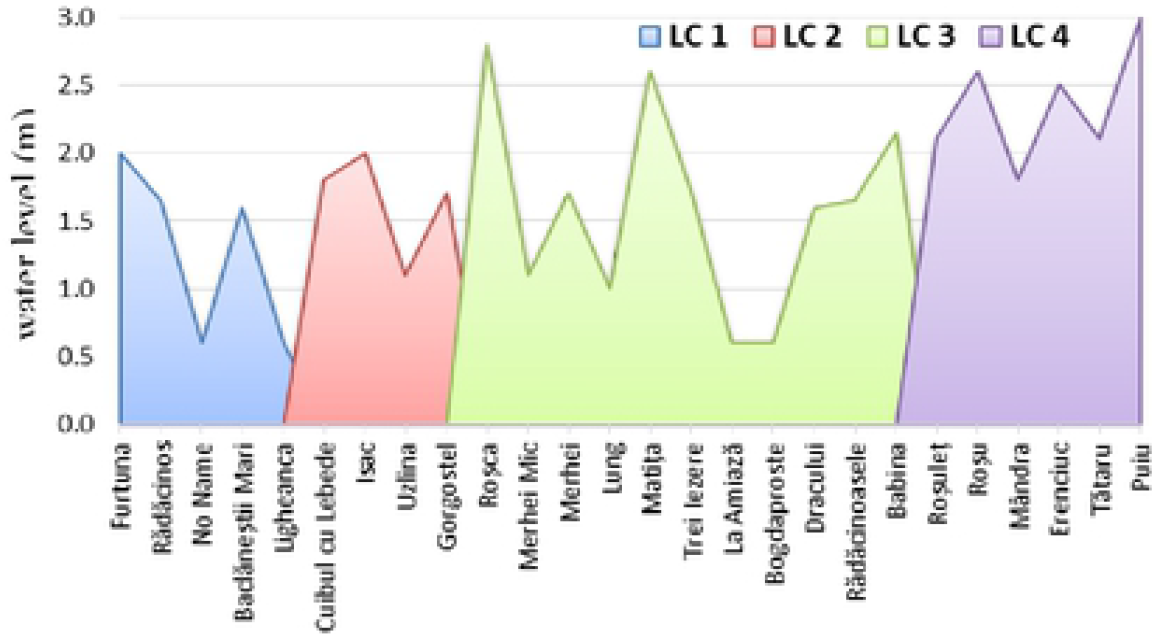
Water level during July 2013 for every studied lake comprised in lake complexes displayed from West to East: Şontea-Furtuna (LC 1), Gorgova - Uzlina (LC 2), Matiţa-Merhei (LC 3), part of the fluvial delta and Roşu-Puiu (LC 4) part of the maritime delta

### 2.2. Water samples collection

Water samples were taken in 2013 from the center of each lake, over the entire water column using a 5 L Schindler – Patalas device. Depending of the lakes depth, the water volume sampled varied between 5 L to 15 L, corresponding to each meter of the lake water layer (1-3 m). From the integrated water sample a variable volume was filtered *in situ* (Table 1) using a vacuum filtration system with 250 mL capacity with a metal hand pomp (Nalgene, USA) and a *glass fibers filter* GF/F, 47 mm, with 0.45 μm pore size (Whatman, UK) until this was saturated with biomass containing environmental DNA (eDNA). Each sample consisted of two GF filters replicates *preserved separately in a zip bag full with silica beads* to be kept dried until the DNA extraction. The same kind of eDNA samples were collected from Dâmboviţa River for performing preliminary extraction tests to prevent from wasting the lake samples.

Another set of eDNA samples were collected in March 2015 from the integrated water samples of the lakes: 30 L were filtered using a phytoplankton *net with 35 μm mesh size* and the obtained biomass was preserved in 200 mL tubes with *ethanol 96%. Fresh water culture* of *Tetradesmus obliquus* (ex *Scenedesmus obliquus*) grown in laboratory for 3-5 weeks on solid BG 11 medium or in WC liquid medium was also used as biomass for DNA extraction. For positive control, *Microcystis aeruginosa* PCC 7806 strain (obtained from the Pasteur Culture collection) was grown in the lab also in WC medium for 3-5 weeks and filtered on GF filters and preserved similar with the eDNA samples.

### 2.3. DNA extraction

For eDNA extraction *Jena Bioscience Animal and Fungi DNA preparation Kit* (JN), *NucleoSpin genomic DNA purification kit* (NS) were used for part of the DDBS lakes samples as well for Dâmboviţa River samples in order to perform preliminary tests. The *xanthogenate nucleic acid isolation method*, based on lab-made extraction buffer called xanthogenate-SDS (XS) was used for all the lakes samples and control strain. For freshwater culture extraction, 10-15 mL of liquid culture were centrifuged to obtain wet biomass, next used for the DNA extraction.The culture grown on solid medium was directly taken from the Petri dish and used for DNA extraction. The XS extraction protocol used was adapted for environmental cyanobacteria from wet biomass preserved in alcohol and GF after Tillett and Neilan (Tillett and Neilan 2000) and is summarized in figure 3 and the recipe is detailed in Table 2 and Table S1. Briefly, 1/8 of each dried GF/F filter was incubated in 750 μL of XS buffer for 3 hours, and then centrifuged for 15 minutes at 22,000 × g. The supernatant was mixed with 750 μL phenol: chloroform: isoamyl alcohol (25:24:1) and centrifuged at 22,000 × g; this process was repeated with the upper aqueous phase. The resulting supernatant was mixed with 1 volume isopropanol and 1/10 volume of 4 M ammonium acetate, kept on ice for 20 minutes and centrifuged at 22,000 × g for 15 minutes. The DNA pellet was washed with 1 mL 70% ethanol, centrifuged for 10 minutes at 22,000 × g, dried for 30 minutes under a sterile hood, and dissolved in 50 μL sterile water. The same protocol was followed for extracting DNA from the reference strain from GF as well as for the filtered with plankton net and centrifuged biomass, and fresh lab cultures of green algae. From the final DNA solution (50 μL) of each sample, two sets of aliquots diluted ten times with sterile MiliQ water were made for avoiding the contamination in the further analysis of the samples. Extracted DNA was quantified by UV spectrophotometry using a NanoDrop ND2000 spectrophotometer (Thermo Fisher Scientific, USA) using 1 μL of 1/10 diluted DNA and for the assesment of the nucelic acid puritiy we measured the optical density (OD) (nm) for the 260/280 and 260/230 absorbance ratio.

**Figure 3.**
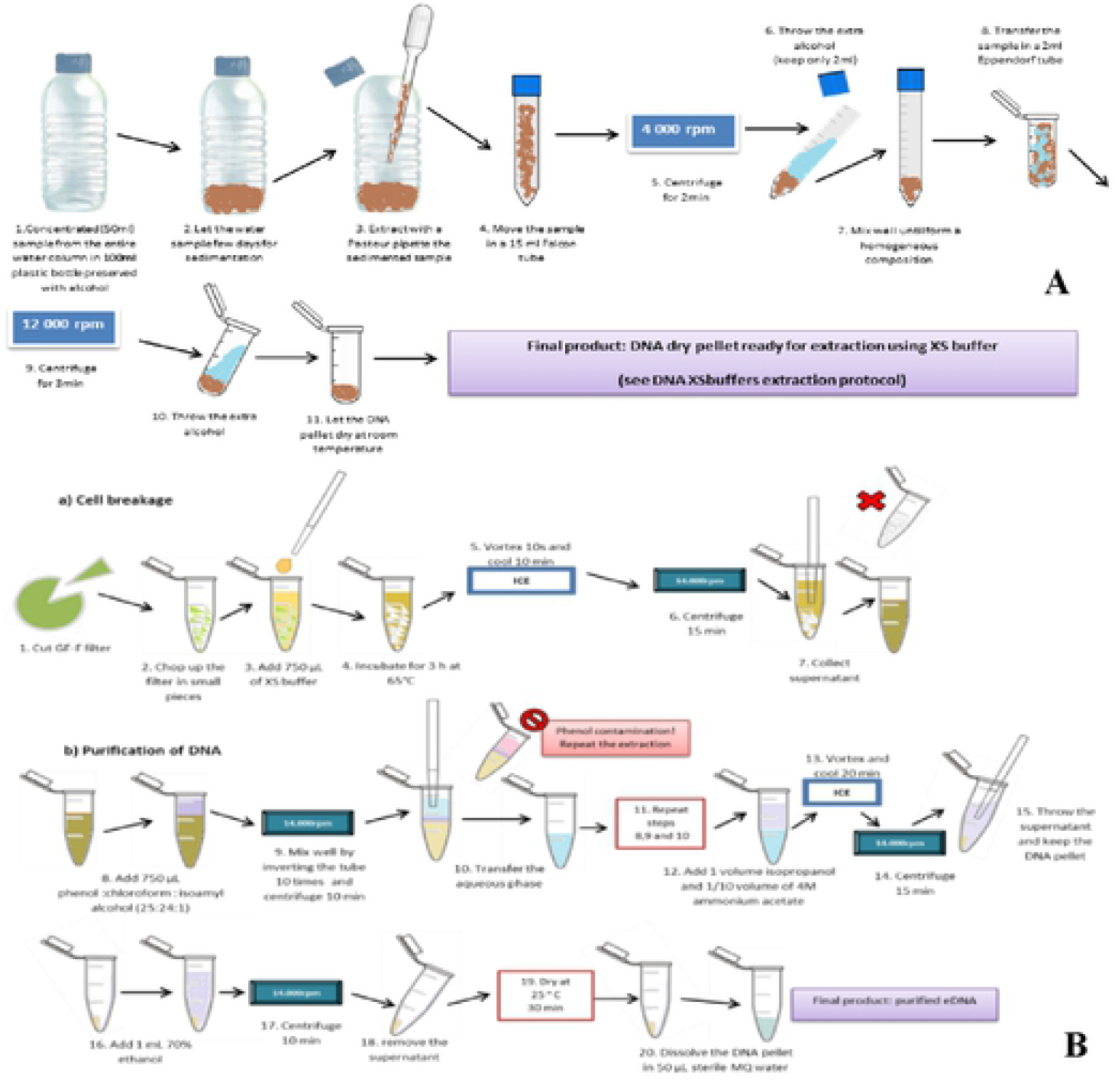
Schematic eDNA extraction protocol using lab-made extraction buffer based on potassium ethyl xanthogenate (XS buffer) for: **A** - biomass preserved in alcohol and **B** - GF filters full with biomass

**Table 2.**
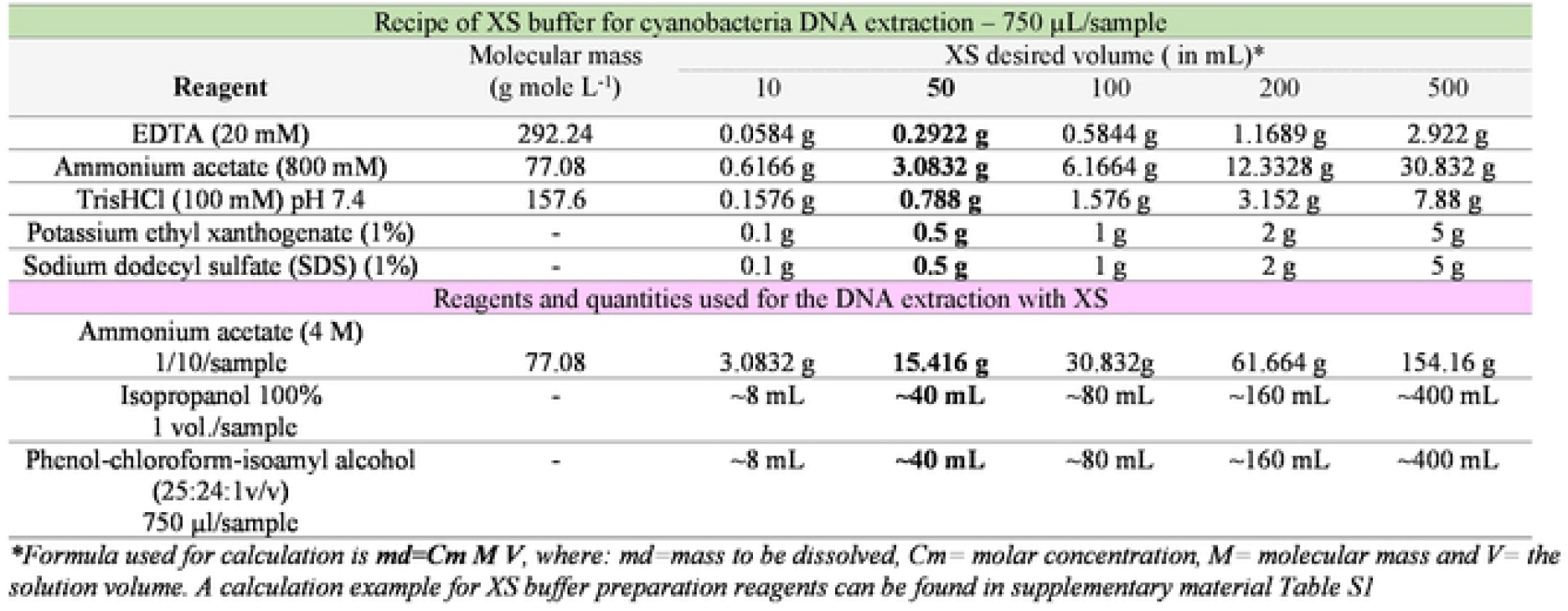
Detailed recipe for reagent preparation for different volume of XS buffer (10-500 mL); for this study each time 50 mL of fresh buffer (bolded column) were prepared; the extraction procedure is detailed in Fig. 3

| Recipe of XS buffer for cyanobacteria DNA extraction – 750 µL/sample |  |  |  |  |  |  |
| --- | --- | --- | --- | --- | --- | --- |
| Reagent | Molecular mass<br>(g mole L<sup>-1</sup>) | XS desired volume ( in mL)* |  |  |  |  |
|  |  | 10 | 50 | 100 | 200 | 500 |
| EDTA (20 mM) | 292.24 | 0.0584 g | **0.2922 g** | 0.5844 g | 1.1689 g | 2.922 g |
| Ammonium acetate (800 mM) | 77.08 | 0.6166 g | **3.0832 g** | 6.1664 g | 12.3328 g | 30.832 g |
| TrisHCl (100 mM) pH 7.4 | 157.6 | 0.1576 g | **0.788 g** | 1.576 g | 3.152 g | 7.88 g |
| Potassium ethyl xanthogenate (1%) | - | 0.1 g | **0.5 g** | 1 g | 2 g | 5 g |
| Sodium dodecyl sulfate (SDS) (1%) | - | 0.1 g | **0.5 g** | 1 g | 2 g | 5 g |
| Reagents and quantities used for the DNA extraction with XS |  |  |  |  |  |  |
| Ammonium acetate (4 M)<br>1/10/sample | 77.08 | 3.0832 g | **15.416 g** | 30.832g | 61.664 g | 154.16 g |
| Isopropanol 100%<br>1 vol./sample | - | ~8 mL | **~40 mL** | ~80 mL | ~160 mL | ~400 mL |
| Phenol-chloroform-isoamyl alcohol<br>(25:24:1v/v)<br>750 µl/sample | - | ~8 mL | **~40 mL** | ~80 mL | ~160 mL | ~400 mL |

\*Formula used for calculation is $md = Cm M V$ , where: $md$ = mass to be dissolved, $Cm$ = molar concentration, $M$ = molecular mass and $V$ = the solution volume. A calculation example for XS buffer preparation reagents can be found in supplementary material Table S1

### 2.4. Molecular assay

The molecular assay was ran on different kind of samples: 1) fresh lab cultures both from liquid and solid medium, 2) filtered on GFs and dried with silica beads, or 3) filtered with phytoplankton net and preserved with alcohol. Before the environmental DNA (eDNA) extraction of DDBR lakes samples, we performed a series of tests in order to find the most suitable method and protocol to isolate as much DNA as possible. The setup consisted in parallel extraction from liquid and solid medium of *T. obliquus* lab culture and of wet (lab culture) and dry (from GF filters) algae biomass. For DNA extraction tests, commercial kits were used for comparison with a special lab-made extraction buffer (XS) (Table 2). In total, five series of tests were performed (Table 3). For PCR reaction primers were selected from literature and different parameters were varied during the assays in order to optimize them. The final PCR protocol was (a) 95°C for 5’, (b) 35 cycles of the following: 95°C for 1’, 60°C for 1’, 72°C for 1’and (c) final elongation step at 72°C for 6’ with pause at 4°C. This protocol had to be changed and adapted depending on some pilot extractions and primers used. For the PCR amplification of cyanobacteria specific 16S rRNA gene, we used forward and reverse recommended primers (Neilan et al. 1997, Saker et al. 2005) with the amplicon size of 782 bp: *27F AGAGTTTGATCCTGGCTCAG* and *809R GCTTCGGCACGGCTCGGGTCGA TA*. For the PCR master mix (MM) *MangoTaq* DNA polymerase and *MyTaq* DNA Polymerase were used and all PCR reagents were from Bioline (London, UK). To reduce the PCR inhibitors 0.4 μg μL^−1^ of bovine serum albumin (BSA) (GeneOn, Ludwigshafen, Germany) was also added. The MM used for cyanobacteria 16S rRNA amplification is presented in the table 3, with the mention that an excel sheet was designed to facilitate the preparation mix according to samples number (Fig. 4). Products were analyzed on 1.5 - 2 % agarose gel with 1× Tris–Borate EDTA (TBE) or 1× Tris base, acetic acid and EDTA (TAE) buffer. We used 5 μL of each amplified DNA stained with 4 μL *ethidium bromide (EtBr*) or 1.25 μL of *pegGREEN* dye for visualization under UV with photo capture system. To mark the wright position of the amplicon the *Gene Ruler 100 bp Plus DNA Ladder* (ThermoFisher Scientific, Waltham, MA, USA) was used with a range band from 100 to 3000 bp (the band of interest being around 800 bp).

**Table 3.** Series of tests performed prior to DNA isolation of DDBR eDNA samples in order to select the best protocol; DNA concentration is in ng μL^−1^

| Test 1: different buffer lysis and matrix (JN) |  |  |  |  |
| --- | --- | --- | --- | --- |
| DNA sample | protocol (a) | 260\280 | protocol (b) | 260\280 |
| TO<sub>liquid</sub>1 | 201.1 | 1.62 | 81 | 1.74 |
| TO<sub>solid</sub> | 16.7 | 1.74 | 13 | 1.37 |
| Test 2: different incubation time and matrix (JN) |  |  |  |  |
| DNA sample | incubation<br>3 hours | 260\280 | incubation<br>over night | 260\280 |
| TO<sub>liquid</sub>2 | 9 | 2.64 | 12.3 | 1.85 |
| DB<sub>filter</sub>1 | 241.2 | 2.11 | 492.7 | 2.08 |
| DB<sub>filter</sub>2 | 163.9 | 2.16 | 335.5 | 2.06 |
| DB<sub>filter</sub>3 | 344.4 | 2.12 | 437.8 | 2.05 |
| DB<sub>filter</sub>4 | 234 | 2.10 | 329.3 | 2.09 |
| DB<sub>filter</sub>5 | 784.6 | 2.01 | 1734.8 | 2.05 |
| DB<sub>filter</sub>6 | 351 | 2.10 | 386.5 | 2.09 |
| Test 3: different incubation time and matrix (NS) |  |  |  |  |
| DNA sample | incubation<br>3 hours | 260\280 | incubation<br>over night | 260\280 |
| TO<sub>liquid</sub>3 | 45.6 | 2.12 | 47 | 1.95 |
| DB<sub>filter</sub>7 | 74.3 | 1.98 | 104.2 | 2.11 |
| LC4PU | 12.5 | 2.01 | 53.8 | 2.02 |
| LC2CL | 8.7 | 1.42 | 14.4 | 1.65 |
| LC3LU | 13.9 | 1.58 | 19.1 | 2.38 |
| Test 4: different extraction kits and matrix (JN versus NS) |  |  |  |  |
| DNA sample | JN | 260\280 | NS | 260\280 |
| TO<sub>liquid</sub>2 | 12.3 | 1.85 | 45.6 | 2.12 |
| DB<sub>filter</sub>8 | 154.9 | 2.11 | 246.35 | 1.98 |
| LC2CL | 3.5 | 2.78 | 14.4 | 1.65 |
| LC3LU | 3.9 | 1.84 | 19.1 | 2.38 |
| Test 5: commercial kits versus lab-made buffer (JN, NS, XS) |  |  |  |  |
| DNA sample | JN | NS | XS | 260\280 |
| DB<sub>filter</sub>8 | 154.9 | 246.35 | 934.5 | 1.63 |
| LC2CL | 3.5 | 14.4 | 41.4 | 1.59 |
| LC3LU | 3.9 | 19.1 | 49.4 | 1.76 |

\*TO=*T. obliquus*, DB=Dâmbovița River, LC2CL=Cuibul cu Lebede lake,
LC3LN=Lung lake, LC4PU=Puiu lake, JN= Jena Bioscience Animal and
Fungi DNA preparation Kit, NS= NucleoSpin genomic DNA purification kit
and XS= xanthogenate nucleic acid isolation method

**Figure 4.**
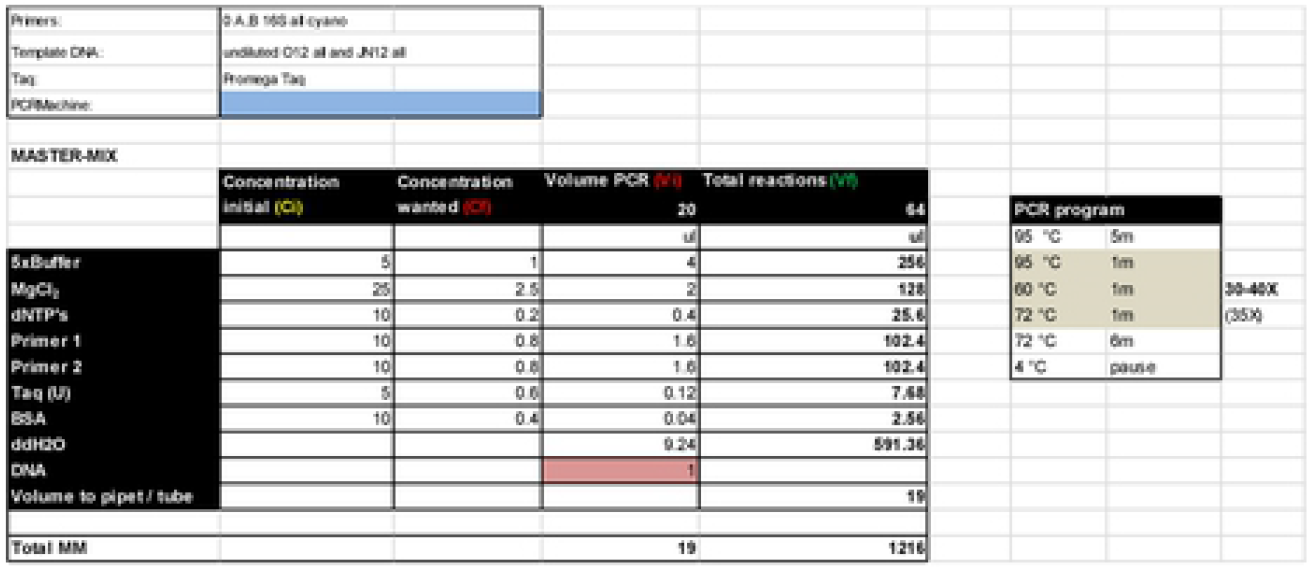
Optimized PCR program and the example of excel recipe calculator for the master mix used for 16S rRNA amplification specific for all cyanobacteria species; for the PCR reaction I μL from the eDNA was used

## 3. Results

### 3.1. DNA isolation preliminary tests

All the tests are described in Table 3 as follows:

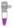 *Test 1: JN protocol (a) and (b) from JN and liquid or solid matrix?* The commercial extraction kit provided by *Jena Bioscience Animal and Fungi DNA preparation Kit* (JN) offers two different types of protocol for extraction: one used also for animal tissues (a) and the second one adapted for fungi (b). For this test fresh algae biomass of *T. obliquus* (TO) cultivated both in WC liquid medium and on BG 11solid medium was used.
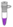 *Test 2: JN short or long time incubation period and dry or wet matrix?* A second round of tests was performed to evaluate the DNA extraction efficiency at different incubation periods, 3 hours compared to overnight (12 hours) from two types of matrix: wet, the same as in test 1, and dried biomass obtained by filtration of water from Dâmboviţa River on GF/F (1/4 of the filter was used for the DNA extraction).
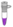 *Test 3: NS short or long time incubation period and dry or wet matrix?* Third round of tests was done as same as test 2 but this time using NS kit for wet and dried biomass as matrix for DNA extraction but also for three of our DDBS eDNA samples (1/8 of GF) chosen from lakes belonging to different lake complexes: lake Puiu from LC4, lake Cuibul cu Lebede from LC2 and lake Lungu from LC3.
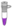 *Test 4: JN versus NS extraction* With this test we compared the two commercial kits in DNA extraction efficiency and purity after over night incubation period. For that, wet and dried biomass were used as matrix for DNA extraction and two of our DDBS eDNA samples (1/8 of GF): lake Cuibul cu Lebede from LC2 and lake Lungu from LC3.
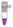 *Test 5: kits versus XS* Last test consisted in the comparison of the commercial kits with the XS buffer from eDNA samples filtered on GF and incubated over night for cells lysis.

#### DNA isolation from DDBR shallow lakes

For all the eDNA samples from DDBR we extracted DNA both with commercial kits and the XS buffer in order to highlight the difference on a larger set of samples (from 26 lakes) collected in different seasons. In the figure 5 the DNA concentration isolated from 1/8 part of GF is exposed comparative for both extraction methods. The quality of the extracted DNA was evaluated measuring the OD express as 280/260 and 260/230 absorbance ratio and the mean values for spring and summer are given in table 4. The total genomic DNA from GF of DDBR shallow lakes over all three seasons is given in figure 6 and for the samples preserved in alcohol (Table S2) the average DNA concentration was 350 ng μL^−1^ (varied between 103 - 793 ng μL^−1^) and OD at 260/280 was 1.64 and at 260/230 was 1.47.

**Fig. 5.**
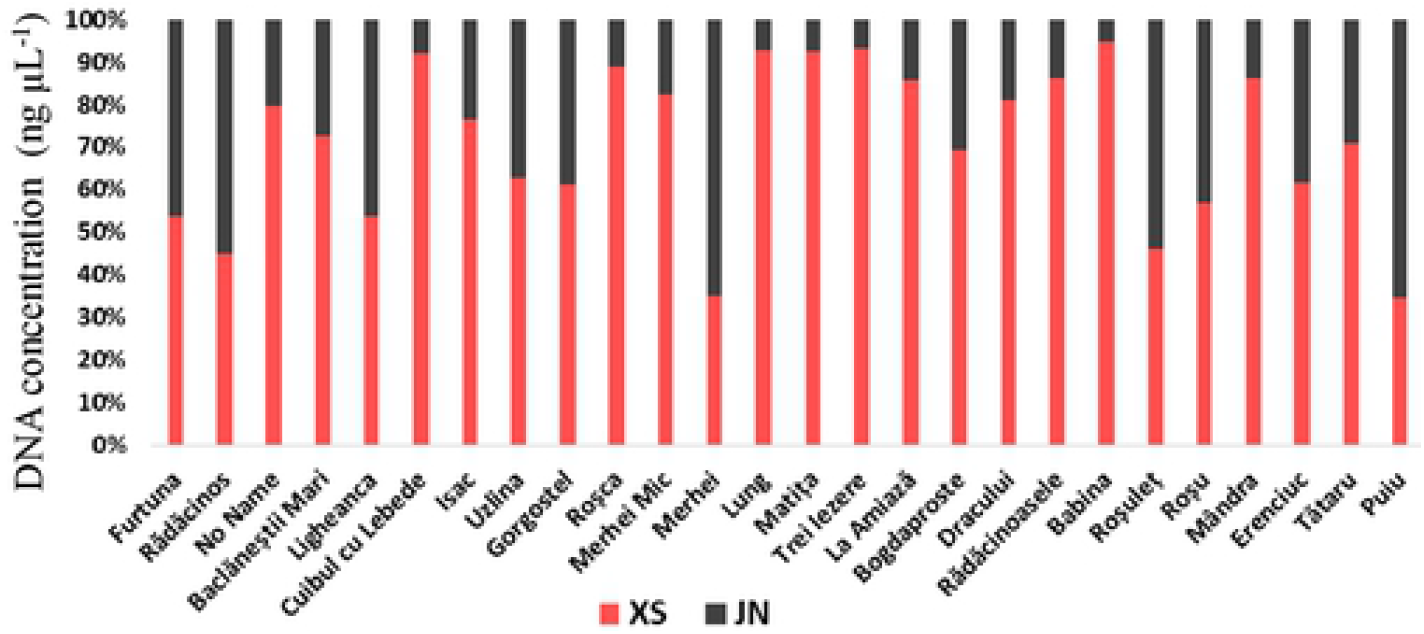
Comparison of totalgenomic DNA from Danube Delta shallow lakes in July 2013 (exception Roşca that was sampled May) extracted with XS buffer (XS) and using Jena Bioscience Animal and Fungi DNA preparation Kit (JN)

**Table 4.** Comparative mean values for the OD ratio from DDBR shallow lakes DNA samples over two seasons obtained with two different extraction methods

| Extraction kit/ season | NanoDrop measurement of OD ratio |  |  |  |
| --- | --- | --- | --- | --- |
|  | 260/280 |  | 260/230 |  |
|  | May | July | May | July |
| JN | 2.045 | 2.0464 | 1.296 | 1.3576 |
| XS | 1.703 | 1.7152 | 2.0065 | 1.8184 |

**Figure 6.**
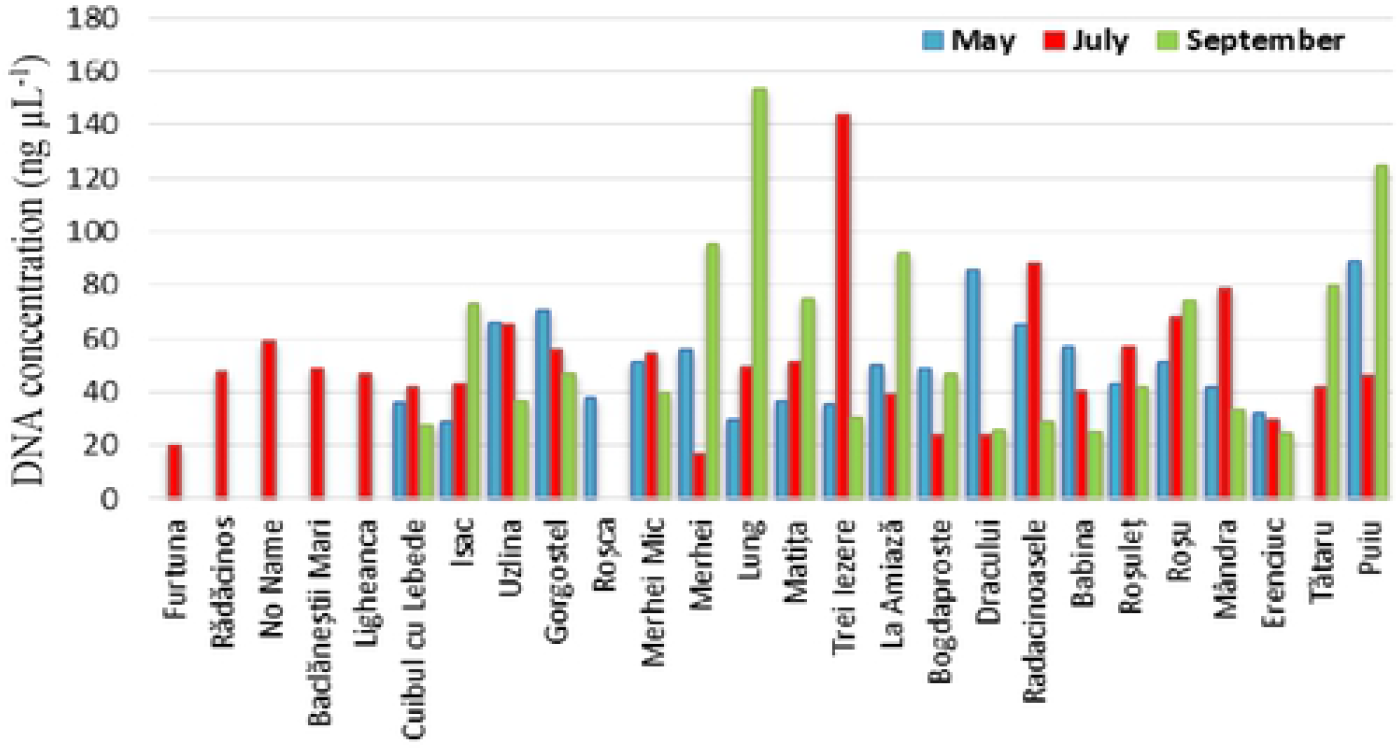
Total genomic DNA from Danube Delta shallow lakes in 2013; for the extraction with XS buffer only 1/8 part of the GF full with biomass was used. Not all lakes were sampled in all three season, from where the missing data

### 3.2. PCR protocol optimization

For PCR optimization we selected and tested the programs and recipes found in the literature (Table 5). The selection criteria consisted in choosing the paper in which scientists used the same primers specific for cyanobacterial 16S rRNA. The annealing temperature was optimized empirically by performing PCRs with 30 to 40 incubation cycles, denaturation and primers anneling at 30’’, 1’ and 1’ and 30’’. To optimize the PCR reaction MM three sets of tests were performed on our DDBS lakes samples as follow:

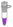 *Test 1* – PCR protocol used was according to the literature: 0.5 mM of MgCl_2_ and 40 cycles (Fig. 7A),
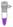 *Test 2* – 2 mM of MgCl_2_ and 40 cycles (Fig. 7B), and
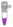 *Test 3* – 2.5 mM of MgCl_2_ and 35 cycles (Fig. 7C).

**Table 5.** Comparative PCR programs and master mix (MM) recipes tested

| Reference/<br>PCR program | (Neilan et al., 1997) | (Saker et al. 2005) | (Jungblut et al. 2005) | Our study |
| --- | --- | --- | --- | --- |
| Initial denaturation | 94°C for 4' | 92 °C for 2' | 92°C for 2' | 95°C for 5' |
| # of cycles | 30 | 35 | 30 | 30-40 |
| Denaturation | 94°C for 20" | 94°C for 10" | 92°C for 20" | 95°C for 30" - 1':30" |
| Primer annealing | 50°C for 30" | 60°C for 20" | 50°C for 30" | 60°C for 30" - 1':30" |
| Strand extension | 72°C for 2' | 72°C for 1' | 72°C for 1' | 72°C for 1' |
| Final extension | unprovided | 72°C for 5' | 72°C for 7' | 72°C for 6' |
| Pause | 4°C | 4°C | 4°C | 4°C |
| Reference/<br>PCR MM | (Neilan et al., 1997) | (Saker et al. 2005) | (Jungblut et al. 2005) | Our study |
| Buffer (μL) | unprovided | unprovided | 1 | 1 |
| MgCl<sub>2</sub> (mM) | unprovided | unprovided | 2.5 | 2.5 |
| dNTP's (mM) | unprovided | unprovided | 0.2 | 0.2 |
| Primers (pM μL<sup>-1</sup>) | unprovided | unprovided | 0.2 | 0.3 |
| Taq (units) | unprovided | unprovided | 0.5 | 1 |
| BSA (μg μL<sup>-1</sup>) | unprovided | unprovided | - | 0.4 |

*\*MM=PCR master mix, dNTP's= deoxyribonucleotide triphosphate, Taq= Taq DNA*
*Polymerase, BSA= bovine serum albumin*

**Figure 7.**
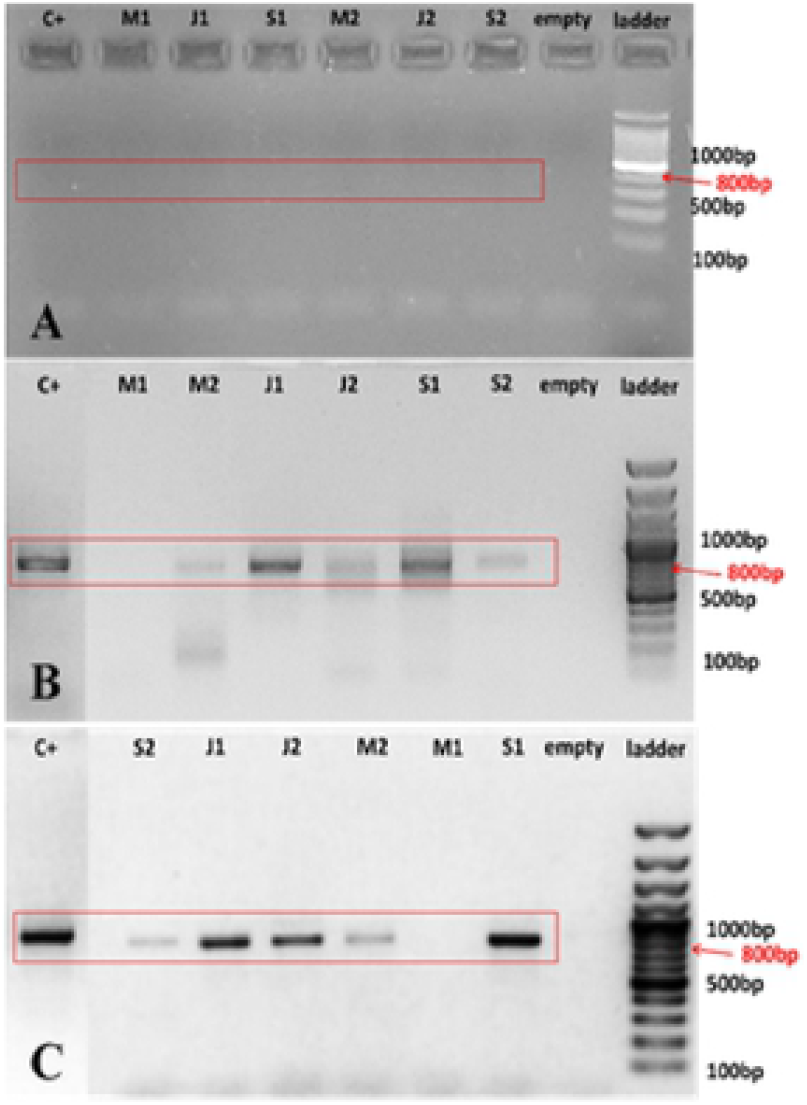
Electrophoresis gels (1.5% agarose, 1 × TAE, dyed with EtBr, ran at 50 V for 40’) used for protocol optimization tests performed with *MyTaq* and 5 μL PCR product (expected amplicon size was 782 bp corresponding to the 16sRNA region specific for cyanobacteria) and using: **A** - 1.5 mM of MgCl_2_ and 30 cycles for PCR program, **B** - 2 mM of MgCl_2_ and 40 cycles for PCR program and **C** - 2.5 mM of MgCl_2_ and 35 cycles for PCR program. Random selection was made from May (M1, 2) July (J1, 2) and September (S1, 2) from eDNA lakes samples. PCR product was amplified in 5 of the 6 samples tested and the intensity of the band varied according to the DNA quantity

After the PCR protocol optimization several tests were ran to verify the presence of 16S rRNA region specific for cyanobacteria in our samples (Fig. 8).

**Figure 8.**
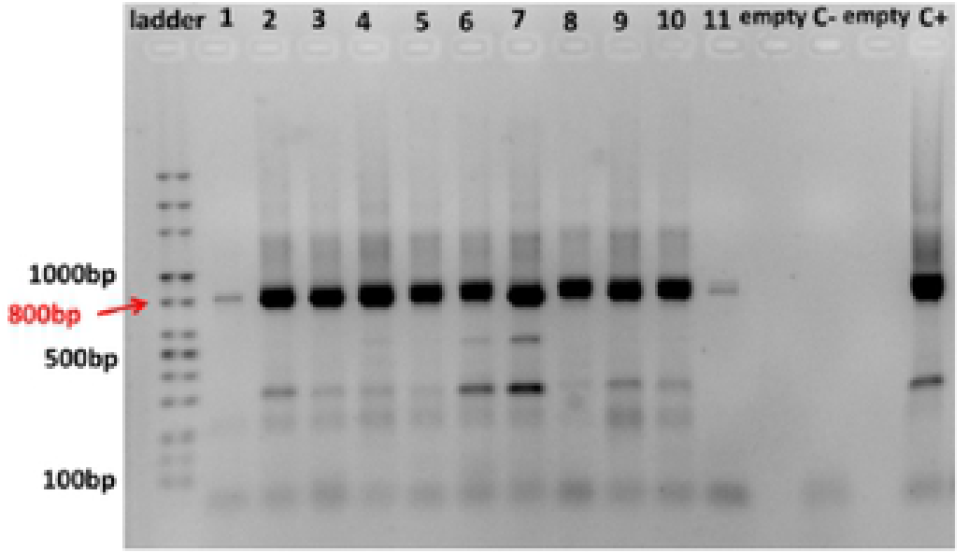
Electrophoresis gel (2 % agarose, 1× TBE buffer, dyed with pegGREEN directly in the gel, ran at 120 V for 50’) performed with *MangoTaq* and 20 μL PCR product (expected amplicon size was 782 bp corresponding to the 16sRNA region specific for cyanobacteria) and using the optimized PCR protocol with BSA (Fig. 7C). Random eDNA from lakes sampled in autumn (1-11) and positive control DNA of *M.aeruginosa* PCC 7806 were used for tests. PCR product was amplified in all the samples, with the mention that in 11 a very thin band was noted meaning less DNA, also unspecific bands are present

## 4. Discussion

### 4.1. Different matrix and DNA extraction kits

Testing the DNA extraction methods is an important and mandatory step no matter what the matrix is, as previous studies showed (Jiang et al., 2005; Gaget et al., 2017), since this represent a critical step in culture-independent bacterial profiling (Hallmaier-Wacker et al. 2018). Even if many different protocols and commercially kits have been developed for DNA extractions from environmental samples (Chandraa et al. 2010) there is still a lack of reliable techniques for DNA extraction as well for RNA isolation for several types of cyanobacteria (Tiam et al. 2019). Almost all commercially kits improved significantly the final purification product for most of bacteria but not for cyanobacteria (Kaczyńska et al. 2013).

Therefore, according to our first hypothesis, we evaluated whether using different biomass matrix and extraction methods the quantity and quality of algae/cyanobacteria extracted DNA from DD could be improved. We showed that biomass extracted from liquid medium instead of biomass obtained from solid culture medium coop better with the extraction buffer specific for tissues provided by the commercial kit from Jena and not the one specific for fungi (Table 3). Since matrix effect was reduced by using the same samples, the variations in the data can be attributed exclusively to the effects of extraction methods as other scientist remarked (Abdel-Latif and Osman 2017). This was an important step since one need to know from which matrix one can extract adequate DNA if proceed further with the cyanobacteria or algae isolation and PCR amplification as well. Preliminary tests were made using a green alga from Pasteur collection in order to minimise the problem of DNA extraction given by the cyanobacteria cell envelope. However, due to the high variability of cyanobacteria shape and size (unicellular, colonial, or filamentous) a variety of cell lysis methods are required (Tiam et al. 2019). Therefore in the second and third set of tests we compared the incubation period necessary for cells lysis and the buffers from two different extraction kits, this time also from eDNA sampled with GF. As expected, more DNA was obtained after overnight incubation period. Also, filtered samples were easier to be long time stored and the DNA quantity was higher since this method allowed to concentrate the biomass from a higher water volume. Further, comparative tests using two different commercial kits revealed that better DNA values were obtained using NucleoSpin kit, especially from our the Danube Delta eDNA samples.

Still, a comparative study in which six DNA extraction kits were tested revealed that PCR inhibitors were present in all DNA solutions extracted (Jiang et al. 2005). Hence, a complete efficient method to obtain higher quantity of DNA and get rid of all the inhibitors in the same time doesn’t exist. However, another serious problem is that DNA isolation and purification efficiencies vary considerably from one species to another (Kaczyńska et al. 2013). Similar to DNA extraction kits, the choice of sample storage buffer has been shown to influence the detected bacterial community (Hallmaier-Wacker et al. 2018). Therefore, ne needs to be careful when using buffers that contain ETDA since this can be found commonly in some elution buffers of commercial kits and in some certain concentrations it may deplete magnesium ions and thus inhibit DNA polymerase activity (Schrader et al. 2012). Also, commercial kits are expensive when a large number of samples need to be processed, while the lab made buffer for DNA extraction represent a more economic extraction method if the efficiency criteria is required.Within this study we tested both several commercial kits and a lab made buffer for the cyanobacteria extraction since it was proven that the extracted DNA concentration varied significantly between the commercial kits (Gaget et al. 2017), which is also in line with our findings (Table 3, test 5).

### 4.2. Commercial DNA extraction kits versus lab-made XS buffer

It is already known that the efficacy of sample processing and DNA extraction may be affected by the most of the known organic compounds e.g.: bile salts, urea, phenol, ethanol, polysaccharides, sodium dodecyl sulphate (SDS), as well as different proteins (Schrader et al. 2012) but the XS protocol has been previously shown to give high-quality genomic DNA, not only from cyanobacteria but from other microorganism as well (Tillett and Neilan 2000). Therefore, in order to test the lab-made buffer for the XS extraction method and in parallel the above mentioned kits, we selected also few samples from the Danube Delta eDNA samples as following: lake Puiu from LC4, considered with high abundance of cyanobacteria, lake Cuibul cu Lebede from LC2 chosen for less cyanobacteria and lake Lungu from LC3 considered with high quantity of inhibitors (Table 3).

It can be easily noticed in the figure 5 that when using the XS extraction method substantial more total genomic DNA was obtained than when the commercial DNA extraction kit Jena Bioscince Animal and Fungi DNA preparation Kit was used. Thus we decided to use the lab made buffer, fresh each time, for the assurance of a high quantity of cyanobacteria DNA for our total genomic DNA samples. Only four out of 26 lakes did not follow the tendency of results, namely lakes Rădăcinos, Merhei, Roşuleţ and Puiu but the eDNA quantity was more than enough to be used for PCR amplification. We cannot explain for sure this unusual result, an explanation could be that during the extraction protocol, part of the DNA pellet didn’t totally dissolved, or, most probable, during the DNA quantification with the NanoDrop we did not manage to mix well the sample, and part of the eDNA remained attached in the bottom part of the tube, undissolved completely. Surprisingly, Lungu lake considered with a high quantity of inhibitors had the highest efficiency with XS method (Fig. 5), therefore we performed all the extraction using this method. However, our DNA pellets after XS isolation were often brownish in colour, suggesting the presence of humics (Yilmaz et al. 2009). Even so, our samples still produce clear band even from smaller DNA fragments in PCR amplification using the universal primers for 16S rDNA gene, therefore this extracted DNA can be successfully applied in different molecular biology methods as other researches experience (Chandraa et al. 2010).

We also found one paper that reported significant lower amount of DNA isolated with the phenol–chloroform method compared with the commercial kit used (Gaget et al. 2017). Contrary, this was not the case with our tests, no matter the matrix that we used for extraction, ten times more DNA values were obtained using this protocol comparing with the commercially kits (Fig.5) and our result was not an isolated case since other scientists experienced similar results when using a phenol-chloroform based extraction method (Palomo et al. 2017). For example, it has been demonstrated that filamentous cyanobacteria respond better to phenol and SDS extraction (Kaczyńska et al. 2013) even if the SDS inhibits the extraction, or that phenol-chloroform extraction is the most efficient in obtaining good quality DNA even from matrix preserved with paraffin (Mirmomeni et al. 2010). For this tests we considered Cuibul cu Lebede as being the lake with less cyanobacteria and Lungu and Puiu with cyanobacteria blooms (Table 3). The best method seems to comprise the filtration of water samples on GF, extraction of DNA (from 1\8 of GF) using XS buffer to obtain a higher amount of DNA comparing with commercial kits (Table 3, Fig. 5), and 1/10 dilution of DNA before PCR (data not shown), as well as the reduction of the amount of primers and utilization of BSA to attenuate the inhibitory effect (Fig. 8). The most probable explanation why XS methods was more efficient is that some strains of cyanobacteria might be harder to lyse than others especially filamentous ones, that are abundant in our samples, and commercial kits extraction columns and buffers are not as efficient as XS. If the cells are not completely lysed (these can easily be observed once the color changes and the extraction buffer becomes more intense), extra vortex and incubation of the samples should be performed until complete cell lysis. One can even add some silica beads and vortex the samples repeatedly and check under microscope to see if the cells lyse or filaments break up, otherwise continue to vortex as long as it is necessary.

### 4.3. DNA quality

In order to establish the quality of each extracted DNA sample we measured it spectrophotometrically using a NanoDrop instrument, since the absorbance profile was useful for detection of contaminants that could severely affect the DNA purity (Abdel-Latif and Osman 2017). Therefore, the A_260_/A_280_ and A_260_/A_230_ ratios for all extractions must be measured to describe the DNA purity beside the quantity (Sellers et al. 2018). Ideally, the pure and undegraded genomic DNA must be characterized by an A_260_/A_280_ ratio of about 1.8, especially in the case of commercial kits and an A_260_/A_230_ ratio of about 2.0 (Kaczyńska et al. 2013).

For our samples (Table 4) the mean value of the A_260_/_280_ ratio was 1.7 which means that our DNA was pure enough, but slightly traces of proteins resulted during the extraction procedure of XS method, while the kit used managed to be protein free. In similar studies for example, the A260/A280 ratio between 1.93 and 2.27 indicates insignificant levels of contamination, while a ratio from 1.6 to 1.8 indicates that the extracted DNA had high purity with absence of proteins and phenols (Abdel-Latif and Osman 2017).

In order to take into consideration also the influence of the local environmental conditions on the eDNA samples, we compared both spring OD values (flooding period with high quantity of sediment/humic acid) with the summer ones, considered mostly with cyanobacteria/alga mass development, therefore with polysaccharides presence. In the case of the XS method, the ratio was nearly 1.7 indicating the sufficient removal of protein contaminants (Chandraa et al. 2010). Even for different species, the DNA extraction efficiency may vary being reflected by the A260/A280 as it has been already shown: 1.84-2.02 for *Anabaena sp*., 1.88-2.12 for *Nodularia spumigena* and 1.75-1.90 for *Nostoc sp*. (Kaczyńska et al. 2013). For the A_260_/A_230_ ratio instead, kits didn’t manage to clean our samples for contaminants either from the environment or from the buffers. On the other hand, with the XS method better results were obtained even if this implied a high risk of phenol contamination, meaning that the genomic DNA extracted by us was suitable for molecular assay.

### 4.4. DNA isolation from DDBR shallow lakes

Since in DDBR is highlighted a strong seasonality regarding the phytoplankton distribution (Moldoveanu et al. 2015), this tendency was found also in the eDNA quantity extracted from our samples during spring, summer and autumn (Fig 6). As expected, higher values were registered both in July and September since according to our data, we found cyanobacteria mass development in some lakes as well in September or even in May (Moza, Postolache et al., 2020). The DNA concentration highlighted this tendency as well, making the XS method a very reliable one for this kind of study. More, in order to be sure that the high quantity of DNA was also purified enough we analyzed the OD ratio mean values for all samples (Table 4). In the case of samples that were preserved with alcohol (Table S2), the A_260_/A_280_ and A_260_/A_230_ ratios were lower, meaning that there was a protein contamination, DNA being more degraded and that the others inhibitors could interfere more with the PCR.

### 4.5. PCR protocol optimization

PCR reaction can fail due to the inhibitors that are very common especially in environmental samples. The PCR inhibitors represent a diverse group of substances such as proteins, salts, and polysaccharides (Abdel-Latif and Osman 2017), that act at different steps of the diagnostic procedure from sampling until the nucleic acids amplification. In our samples from Danube Delta shallow lakes we expected to find the widely occurring freshwater environmental inhibitors represented by: fulvic acids from dead biomass and sediments that copurify with DNA and inhibit PCR and restriction digestion of DNA (Tiam et al. 2019, Yilmaz et al. 2009) (Schrader et al. 2012), humic acids that inhibit PCR and interact with the polymerase preventing the enzymatic reaction even at low concentrations (Kreader 1996, Schrader et al. 2012) (Tiam et al. 2019), as well as the cyanobacterial polysaccharides that may disturb the enzymatic process (Schrader et al. 2012). Calcium salts represent example of inorganic substances with inhibitory effects on the PCR and polymerase activity and even the wall of the reaction tubes (Schrader et al. 2012), powder from gloves (Demeke and Jenkins 2010) or different other salts (e.g. sodium chloride or potassium chloride), detergents and EDTA (Opel et al. 2010) may affect the efficacy of sample processing. Therefore is crucial to select an appropriate extraction method to minimize the inhibitions during samples processing.

In order to annihilate the inhibitors effect in the samples processed by us, especially those from humic acids we added in our PCR mix bovine serum albumin (BSA) in concentration of 400 ng mL^−1^ as other studies recommended (Kreader 1996) (Jiang et al. 2005). Even if this additive is used to boost PCR, is not effective in case of SDS, EDTA and calcium presence (Opel et al. 2010) and in our samples we had them all. BSA, like DMSO is recommended as well for difficult template GC-rich (>60%) like our samples, in order to improve its availability for hybridization and reduces nonspecific binding (Farell and Alexandre 2012).

It was also demonstrated that the bacterial DNA contamination in the Taq polymerase exists and could often give false positive results especially when working with 16S primers (Lupan et al. 2013). This is due mostly to the laboratory environment during the protein purification, therefore we can exclude this possibility in our case since we used specific primers for cyanobacteria. Another potential cause for the bias in the analysis is that the use of 16S primers may favour certain bacterial strains (Hallmaier-Wacker et al. 2018), but in our fist assay attempt we didn’t find any bands, hence the problem was elsewhere.

Nevertheless, we need to take in consideration that the high presence of calcium ions can give a competitive binding to the template and inhibit the PCR reaction, but it might be compensated by the addition of magnesium ions into the MM (Schrader et al. 2012). In order to reduce the inhibitory influence we also diluted the DNA and prepared 1/10 aliquots from the initial eDNA since this method was proven to be widely applied for the dilution of PCR inhibitors (Scipioni et al. 2008). Further, the initial PCR programs and master mix recipes found in others papers were tested and optimized for our DDBR eDNA samples (Table 5). First, the PCR program was modified starting from the one proposed by Neilan and his team in 1997 (Neilan et al. 1997). In order to improve the amplification and to obtain clearer bands, we added more cycles to the initial PCR protocol since other scientists proceeded the same to improve the amplification of low DNA concentration samples in case or 16S rRNA gene amplification (Hallmaier-Wacker et al. 2018). Thus, when assayed the same protocol at 35 and 40 cycles (Fig. 7) clear bands were obtained at 35 cycles (Fig. 7C) compared to 30 cycles (Fig. 7A), while 40 cycle were proven to be too much (Fig. 7B). Results were maintained for diluted DNA samples as well.

However, the annealing of the primers to the DNA template may be disturbed by certain PCR inhibitors (Abbaszadegan et al. 1999) therefore we tested also the diluted aliquots. For example, calcium, mostly resulted from the high diatoms presence in our samples, competed as a cofactor instead of MgCl_2_ binding to our template, resulting in zero amplification for our samples (Fig. 7A). In order to compensate, we increased gradually the MgCl_2_ concentration during our tests, from 1.5 to 2.5 mM (Fig. 7B and C), with the recommendation that the maximum concentration is the most useful.

Other studies also tested different agarose gels concentration (1, 1.5 and 2%) and the best results were observed at the lowest concentration (Abdel-Latif and Osman 2017) which was in concordance with our results. Also, the addition of too much cycles to the PCR program was proven to be not efficient for the amplification since this create chimeric structures as can been seen in the figure 7B. This supports the assumption that lower numbers of cycles are favourable for amplicon sequencing (Hallmaier-Wacker et al. 2018). All the tested samples were amplified with the optimized protocols, even those with a very low DNA quantity of cyanobacteria (Fig. 8), thus the protocols were useful for the analysis of cyanobacteria from DDBR.

## 5. Conclusions and recommendations

For studying freshwater cyanobacteria, especially from Danube Delta Biosphere Reserve shallow lakes, the best eDNA sampling method is using GF saturated with biomass, dried with silica beads and stored for the long term period in the lab. Our recommendation is to use only a small part of the filter full with eDNA since according to our tests, 1/8 of the filter was enough. The best extraction method was proven to be the lab-made buffer based on potassium ethyl xanthogenate (XS). It should be always prepared in small quantities according to the samples number (10-50 mL) since is mandatory to be prepared fresh, in the same day or with a day before, in order to ensure a high quantity and quality of the extracted DNA. The method must be adapted for each sample apart (meaning that the aqueous volume may differ) to avoid phenol contamination, depending on DNA quantity which may vary from lakes to lakes but mostly seasonally: in spring we had less DNA and in summer considerable more, for instance. This means that not all the time the isopropanol volume that need to be added will be the same for all the samples in the same extraction. We strongly recommend to pay attention on this during extraction process. For liquid samples, at least 15 L of water must be filtered through a phytoplankton net to obtain enough biomass for DNA extraction, if there is no possibility to use *in situ* a filtering system with a GF of maximum 45 μm pore size. We also recommend to dilute the extracted DNA ten times and prepare at least two sets of aliquots and to use only a small amount of primers (below 0.3 μL/sample) for PCR reaction, since we had primers residues. Also, never use less than 1 unit of Taq per sample as a general rule. The PCR parameters must be set different as well; the best protocol is to use 35 cycles and vary the MgCl_2_ concentration when is necessary, but not lower than 2% and add BSA since the inhibitors are present in high percent in shallow lakes. Method optimization was an essential step, the findings showing that a combination of various working protocols and reagents is needed to ensure obtaining of as large DNA quantities (and its quality) as possible. Work on further DNA purification, creation of clone library and lakes classification based on occurrence of genes with potential to release toxins is in progress.

## Acknowledgments

This work was supported by the Swiss Enlargement Contribution (IZERZ0 – 142165, “CyanoArchive”) in the framework of the Romanian-Swiss Research Programme. The authors transmit their special thanks to CyanoArchive Project team and Sulina field team for all the support during the sampling trips, to dr. Piet Spaak, dr. Francesco Pomati, dr. Beatrice Kelemen, acad. dr. Octavian Popescu and dr. Iulia Lupan for all the help and supervising, to dr. Marie Eve Monchamp, dr. Christoph Tellenbach, dr. Bogdan Drugă, as well to Esther Keller and Pravin Ganesanandamoorthy for helpful tips in the lab. We also thank dr. Liliana Török for proving us some samples. Special thanks goes to dr. Adrian Florea and to dr. Carmen Chifiriuc for the scientific input and text correction. Authors also acknowledge the specialists from Genetic Diversity Centre of ETH Zürich for their helpful discussions and advices. The authors are grateful to the reviewers and Journal Editorial board.

## Author contribution

MIM designed and performed the experiments, CP supervised and validated the results, and both authors wrote, discussed the results and commented on the manuscript.

## Tables

for

Moza and Postolache, 2021, *Journal Plos One*

## References

Abbaszadegan, M., Stewart, P. and LeChevallier, M. (1999) A strategy for detection of viruses in groundwater by PCR. Applied and environmental microbiology 65(2), 444–449.

Abdel-Latif, A. and Osman, G. (2017) Comparison of three genomic DNA extraction methods to obtain high DNA quality from maize. Plant Methods 13(1), 1.

Akkak, A., Boccacci, P. and Botta, R. (2008) An efficient dna-extraction protocol for nut seeds. Journal of food quality 31(4), 549–557.

Albrecht, M., Pröschold, T. and Schumann, R. (2017) Identification of Cyanobacteria in a Eutrophic Coastal Lagoon on the Southern Baltic Coast. Frontiers in microbiology 8(923).

Billi, D., Caiola, M.G., Paolozzi, L. and Ghelardini, P. (1998) A method for DNA extraction from the desert cyanobacterium Chroococcidiopsis and its application to identification of ftsZ. Appl. Environ. Microbiol. 64(10), 4053–4056.

Chandraa, S., Varshneya, A.K., Sinhaa, S. and Mathurb, N. (2010) A Rapid and Efficient DNA Extraction Method for PCR Based Assays in Activated Sludge. Jour Pl Sci Res 26(2), 213–218.

Coops, H., Buijse, L.L., Buijse, A.D.T., Constantinescu, A., Covaliov, S., Hanganu, J., Ibelings, B.W., Menting, G., Navodaru, I. and Oosterberg, W. (2008) Trophic gradients in a large-river Delta: ecological structure determined by connectivity gradients in the Danube Delta (Romania). River Research and Applications 24(5), 698–709.

Demeke, T. and Jenkins, G.R. (2010) Influence of DNA extraction methods, PCR inhibitors and quantification methods on real-time PCR assay of biotechnology-derived traits. Anal Bioanal Chem 396(6), 1977–1990.

Dobhal, S., Zhang, G., Rohla, C., Smith, M.W. and Ma, L.M. (2014) A simple, rapid, cost-effective and sensitive method for detection of Salmonella in environmental and pecan samples. J Appl Microbiol 117(4), 1181–1190.

Drábková, L., Kirschner, J. and Vlĉek, Ĉ. (2002) Comparison of seven DNA extraction and amplification protocols in historical herbarium specimens of Juncaceae. Plant Molecular Biology Reporter 20(2), 161–175.

Enache, I., Florescu, L.I., Moldoveanu, M., Moza, M.I., Parpală, L., Sandu, C., Turko, P., Rîşnoveanu, G. and Spaak, P. (2019) Diversity and distribution of Daphnia across space and time in Danube Delta lakes explained by food quality and abundance. Hydrobiologia 842(1), 39–54.

Farell, E.M. and Alexandre, G. (2012) Bovine serum albumin further enhances the effects of organic solvents on increased yield of polymerase chain reaction of GC-rich templates. BMC research notes 5(1), 257.

Fontana, S., Thomas, M.K., Moldoveanu, M., Spaak, P. and Pomati, F. (2018) Individual-level trait diversity predicts phytoplankton community properties better than species richness or evenness. The ISME Journal 12(2), 356–366.

Gaget, V., Keulen, A., Lau, M., Monis, P. and Brookes, J. (2017) DNA extraction from benthic Cyanobacteria: comparative assessment and optimization. Journal of Applied Microbiology 122(1), 294–304.

Hallmaier-Wacker, L.K., Lueert, S., Roos, C. and Knauf, S. (2018) The impact of storage buffer, DNA extraction method, and polymerase on microbial analysis. Scientific reports 8(1), 6292.

Irimuş, I.A. (2006) The hydrological regime of the Danube River in the deltaic sector. In: Tudorancea C, Tudorancea MM, editors. Danube Delta - Genesis and Biodiversity. Leiden, The Netherlands: Blackhuys Publishers, 53–64.

Jiang, J., Alderisio, K.A., Singh, A. and Xiao, L. (2005) Development of procedures for direct extraction of Cryptosporidium DNA from water concentrates and for relief of PCR inhibitors. Appl Environ Microbiol 71(3), 1135–1141.

Kaczyńska, A., Łoś, M. and Węgrzyn, G. (2013) An improved method for efficient isolation and purification of genomic DNA from filamentous cyanobacteria belonging to genera Anabaena, Nodularia and Nostoc. Oceanological and Hydrobiological Studies 42(1), 8–13.

Kreader, C.A. (1996) Relief of amplification inhibition in PCR with bovine serum albumin or T4 gene 32 protein. Appl. Environ. Microbiol. 62(3), 1102–1106.

Lupan, I., Ianc, B.M., Ochiş, C. and Popescu, O. (2013) The evidence of contaminant bacterial DNA in several commercial Taq polymerases. Biotechnol Lett 18, 8007–8012.

Martins, J., Peixe, L. and Vasconcelos, V.M. (2011) Unraveling cyanobacteria ecology in wastewater treatment plants (WWTP). Microbial ecology 62(2), 241–256.

Mirmomeni, M., Sajjadi Majd, S., Sisakhtnezhad, S. and Doranegard, F. (2010) Comparison of the three methods for DNA extraction from paraffin-embedded tissues. Journal of Biological Sciences 10(3), 261–266.

Moldoveanu, M., Zinevici, V., Parpală, L., Ionică, D., Păceşilă, I., Dumitrache, A., Sandu, C. and Florescu, L. (2015) The role of plankton communities in the functional capacity of the Danube Delta ecosystems - a long term study. Muzeul Olteniei Craiova. Muzeul Olteniei Craiova. Studii şi comunicări - Ştiinţele Naturii 31(2), 183–188.

Moore, L.R., Huang, T., Ostrowski, M., Mazard, S., Kumar, S.S., Gamage, H.K.A.H., Brown, M.V., Messer, L.F., Seymour, J.R. and Paulsen, I.T. (2019) Unicellular Cyanobacteria Are Important Components of Phytoplankton Communities in Australia’s Northern Oceanic Ecoregions. Frontiers in microbiology 9(3356).

Moreira, C., Vasconcelos, V. and Antunes, A. (2013) Phylogeny and biogeography of cyanobacteria and their produced toxins. Marine drugs 11(11), 4350–4369.

Morin, N., Vallaeys, T., Hendrickx, L., Natalie, L. and Wilmotte, A. (2010) An efficient DNA isolation protocol for filamentous cyanobacteria of the genus Arthrospira. Journal of Microbiological Methods 80(2), 148–154.

Moza, M.I., Postolache, C., Benedek, A.M., Moldoveanu, M. and Spaak, P. (2021) Geographical and temporal patterns of cyanobacterial assemblages in the Danube Delta lake complexes. Hydrobiologia 848(4), 753–771.

Nair, H.P., Vincent, H. and Bhat, S.G. (2014) Evaluation of five in situ lysis protocols for PCR amenable metagenomic DNA from mangrove soils. Biotechnol Rep (Amst) 4, 134–138.

Neilan, B.A., Jacobs, D., Blackall, L.L., Hawkins, P.R., Cox, P.T. and Goodman, A.E. (1997) rRNA sequences and evolutionary relationships among toxic and nontoxic cyanobacteria of the genus Microcystis. International Journal of Systematic and Evolutionary Microbiology 47(3), 693–697.

Oosterberg, W., Staras, M., Bogdan, L., Buijse, A., Constantinescu, A., Coops, H., Hanganu, J., Ibelings, B.W., Menting, G. and Navodaru, I. (2000) Ecological gradients in the Danube Delta lakes: present state and man-induced changes. Lelystad: Institute for Inland Water Management and Waste Water Treatment RIZA 170.

Opel, K.L., Chung, D. and McCord, B.R. (2010) A study of PCR inhibition mechanisms using real time PCR. J Forensic Sci 55(1), 25–33.

Paerl, H.W. and Huisman, J. (2008) Blooms like it hot. Science 320(5872), 57–58.

Palomo, F.S., Rivero, M.G.C., Quiles, M.G., Pinto, F.P., Machado, A.M.d.O. and Carlos Campos Pignatari, A. (2017) Comparison of DNA Extraction Protocols and Molecular Targets to Diagnose Tuberculous Meningitis. Tuberculosis Research and Treatment 2017, 5089046.

Romanescu, G. (2005) Morpho-hydrographical evolution of the Danube Delta, Editura Pim.

Saker, M., Fastner, J., Dittmann, E., Christiansen, G. and Vasconcelos, V. (2005) Variation between strains of the cyanobacterium Microcystis aeruginosa isolated from a Portuguese river. Journal of Applied Microbiology 99(4), 749–757.

Schrader, C., Schielke, A., Ellerbroek, L. and Johne, R. (2012) PCR inhibitors–occurrence, properties and removal. Journal of Applied Microbiology 113(5), 1014–1026.

Scipioni, A., Bourgot, I., Mauroy, A., Ziant, D., Saegerman, C., Daube, G. and Thiry, E. (2008) Detection and quantification of human and bovine noroviruses by a TaqMan RT-PCR assay with a control for inhibition. Mol Cell Probes 22(4), 215–222.

Sellers, G.S., Di Muri, C., Gómez, A. and Hänfling, B. (2018) Mu-DNA: a modular universal DNA extraction method adaptable for a wide range of sample types. Metabarcoding and Metagenomics 2.

Tiam, S.K., Comte, K., Dalle, C., Duval, C., Pancrace, C., Gugger, M., Marie, B., Yéprémian, C. and Bernard, C. (2019) Development of a new extraction method based on high-intensity ultra-sonication to study RNA regulation of the filamentous cyanobacteria Planktothrix. PloS one 14(9).

Tillett, D. and Neilan, B.A. (2000) Xanthogenate nucleic acid isolation from cultured and environmental cyanobacteria. Journal of Phycology 36(1), 251–258.

Török, L. (2005) Seasonal succesion of phytoplankton from lakes of the Danube Delta. Acta Oecologica XII(1-2), 15–23.

Verbylaite, R., Beisys, P., Rimas, V. and Kuusiene, S. (2010) Comparison of ten DNA extraction protocols from wood of European Aspen (Populus tremula L.). Baltic Forestry 16(1), 30.

Whitton, B.A. and Potts, M. (2007) The ecology of cyanobacteria: their diversity in time and space, Springer Science & Business Media.

Yilmaz, M., Phlips, E.J. and Tillett, D. (2009) Improved methods for the isolation of cyanobacterial dna from environmental samples. Journal of Phycology 45(2), 517–521.

## References

Jungblut, A.D., Hawes, I., Mountfort, D., Hitzfeld, B., Dietrich, D.R., Burns, B.P. and Neilan, B.A. (2005) Diversity within cyanobacterial mat communities in variable salinity meltwater ponds of McMurdo Ice Shelf, Antarctica. Environmental microbiology 7(4), 519–529.

